# Kernel-smoothed permutation for extreme P-value estimation in genetic association studies

**DOI:** 10.1101/2024.01.09.574752

**Authors:** Jiayi Bian, Caifeng Li, Jingjing Wu, Quan Long

**Affiliations:** Department of Mathematics and Statistics, University of Calgary, Calgary, Alberta, Canada; Department of Biochemistry and Molecular Biology, University of Calgary, Calgary, Alberta, Canada; Mathison Centre for Mental Health Research and Education, Hotchkiss Brain Institute, University of Calgary, Calgary, Alberta, Canada; Department of Medical Genetics, University of Calgary, Calgary, Alberta, Canada; Alberta Children’s Hospital Research Institute, University of Calgary, Calgary, Alberta, Canada

**Keywords:** P-value estimation, test statistic, permutation test, kernel-based density estimation

## Abstract

In genetic studies, permutation tests serve as a cornerstone to estimate P-values. This is because researchers may design new test statistics without a known closed-form distribution, or the assumption of a well-established test may not hold. However, permutation tests require vast number of permutations which is proportional to the magnitude of the actual P-values. When it comes to genome-wide association studies where multiple-test corrections are routinely conducted, the actual P-values are extremely small, requiring a daunting number of permutations that may be beyond the available computational resources. Existing models that reduce the required number of permutations all assume a specific format of the test statistic to exploit its specific statistical properties. We propose Kernel-smoothed permutation which is a model-free method universally applicable to any statistic. Our tool forms the null distribution of test statistics using a kurtosis-driven transformation, followed by a kernel-based density estimation (KDE). We compared our Kernel-smoothed permutation to Naïve permutation using statistics from known closed-form null distributions. Based on three frequently used test statistics in association studies, i.e., t-test, sequence kernel association test (SKAT), and chi-squared test, we demonstrated that our model reduced the required number of permutations by a magnitude with the same or higher accuracy.

## INTRODUCTION

In the Hypothesis test, one needs to compute the P-value out of the test statistic using the null distribution (Banerjee et al., 2009). Although this can be done analytically, for many complex situations, especially in genetic studies, the closed form of null distribution may not be available, or the test assumptions are not satisfied. In these scenarios, the permutation test is the default procedure to form the null distribution of the focal statistic (Berry et al., 2011; Chung & Romano, 2013). Permutation tests have desirable properties that can guarantee exactness if data are exchangeable, and the procedure is very straightforward, so it can be adaptable to most test statistics. Consequently, they have become a popular tool among researchers in genetics and other life sciences since they facilitate the design of novel statistics to ascertain statistical significance (Doerge & Churchill, 1996; Nichols & Holmes, 2002; Simpson et al., 2013). Nonetheless, it’s important to recognize the drawbacks of permutation tests. They can be computationally expensive and time-consuming, rendering them impractical to generate all possible permutations in some scenarios.

In genetic studies involving large-scale omics data, the threshold for statistical significance is often set at very small P-value levels due to the multiple testing corrections (Benjamini & Hochberg, 1995). For instance, to account for multiple testing in genome-wide association studies (GWAS), a stringent P-value threshold of 5 × 10^−8^ is commonly adopted to identify the association between a common genetic variant and a trait of interest (Belmont et al., 2005; Pe’er et al., 2008). Likewise, gene-based analyses employ the Bonferroni correction to adjust the P-value threshold, typically calculated as 0.05 divided by the number of genes being tested (Cui & Churchill, 2003; McDermaid et al., 2019). Given these standards, permutation tests can quickly become computationally demanding, especially when estimating exceedingly small P-values across a multitude of tests. Consequently, there is a pressing need for more efficient methods in such high-dimensional settings. Several recent studies have developed efficient algorithms based on permutation tests: Zhang et al. estimated genome-wide significance based on Poisson de-clumping heuristics (Zhang & Liu, 2011); Segal et al. employed fast asymptotic approximation of small P-values by a partitioning and resampling scheme (Segal et al., 2018); and Yang et al. approximated small P-values using sequential Monte Carlo and the Edgeworth expansion (Yang et al., 2019). Despite these advancements, the scope of these methods remains confined to specific test statistic families. Our goal is to bridge this gap by introducing a versatile model-free protocol that applies to any test statistic without limiting its form.

The idea of this work is to replace the generation of vast numbers of permuted test statistics, by maximizing the utility of limited permutated statistics. Kernel-based density estimation (KDE) provides one such approach. KDE serves as a non-parametric way to estimate the probability density function (PDF) of a random variable (Chen, 2017; Jones & Kappenman, 1992; Scott et al., 1917; Silverman, n.d.). Even though KDE’s foundations trace back to the pioneering work of Rosenblatt and Parzen over fifty years ago (Rosenblatt, 1971), continuous advancements in computing have expanded its application across a wide spectrum of scientific fields (King et al., 2016; Tsai & Chen, 2007; Węglarczyk, 2018). By applying a kernel function—like the Gaussian— to each data point and then aggregating them, KDE captures the inherent PDF of the data. This results in a refined and smooth curve that can estimate the probability of any value within the dataset. In KDE, the choice of the kernel function, represented as *K*, and the bandwidth, denoted by *h*, control the smoothness of a density estimate. Notably, when employing KDE to model permuted test statistics, setting the bandwidth to zero yields results parallel to a conventional Naïve permutation test (Epstein et al., 2012).

In our study, we introduce the Kernel-smoothed permutation method, leveraging KDE to model the null distribution of permuted test statistics, with the objective of refining the accuracy of small P-value estimations (e.g., <10^−6^). The integration of a kernel function in KDE enables a broader representation of each permuted point in forming the null distribution’s properties, reducing the number of permutations required. In scenarios with heavy-tailed test statistics that can hamper the efficacy of kernel smoothing, we developed a kurtosis-driven Box-Cox transformation, a step prior to the KDE application. Crucially, as a non-parametric method, Kernel-smoothed permutation does not require restrictive assumptions about parametric families. This flexibility permits its application across all spectrums of test statistics to construct a density that truly captures the inherent structure of the data.

To test the Kernel-smoothed permutation method, we applied it to three test statistics that we know the closed-form distributions (as the gold standard for comparison): a two-sample t-test, the gene-based sequence kernel association test (SKAT) (Wu et al., 2010, 2011) and the chi-squared test. The Wellcome Trust Case Control Consortium (WTCCC) (Burton et al., 2007) data are used for comparison. Through a side-by-side comparison of P-values derived from both Kernel-smoothed permutation and Naïve permutation, we demonstrated that our Kernel-smoothed permutation approach consistently yielded more accurate results than its Naïve counterpart when given the same numbers of permuted test statistics. Notably, with a reduced (10%) set of test statistics of a magnitude, Kernel-smoothed permutation can still outperform Naïve permutation.

In the rest of this paper, we began by providing a comprehensive overview of Kernel-smoothed permutation’s underlying mechanics and rationales, followed by a detailed narrative on methods behind our implementation of the Kernel-smoothed permutation technique. At the end, a presentation of our results from the application of this method to the trio of above-mentioned tests and a discussion of potential avenues for further research is provided.

## MATERIALS AND METHODS

### The conceptual framework underlying the Kernel-smoothed permutation

In the Naïve permutation, each permuted sample is self-representative. After generating a substantial set of these permuted samples, they are arranged in ascending order, allowing us to determine the rank of the test statistic (in the original hypothesis test) among these samples (**Figure 1a**). Conversely, the Kernel-smoothed permutation allows each permuted sample to represent a broader region in the format of a kernel-based distribution (illustrated by the contours surrounding each sample in **Figure 1b**). Here the kernel function *K* determines their shapes and the window *h* determines their bandwidth. The kernel-estimated null distribution is formed by aggregating these sample-centered distributions, leveraging extra information covered by the bandwidth of these kernels. Such a kernel-smoothing can reduce the number of permutations required by exploiting each sample more thoroughly than a Naïve permutation (which is equivalent to a kernel with zero bandwidth). In genetic association studies, when computing a very small P-value, the heavy-tailed distribution can undermine the efficacy of kernel smoothing, which mandates a transformation of samples. We have then developed a kurtosis-driven Box-Cox transformation that narrows down the relative excess kurtosis towards a normal distribution (**Figure 1c**).

**Figure 1.**
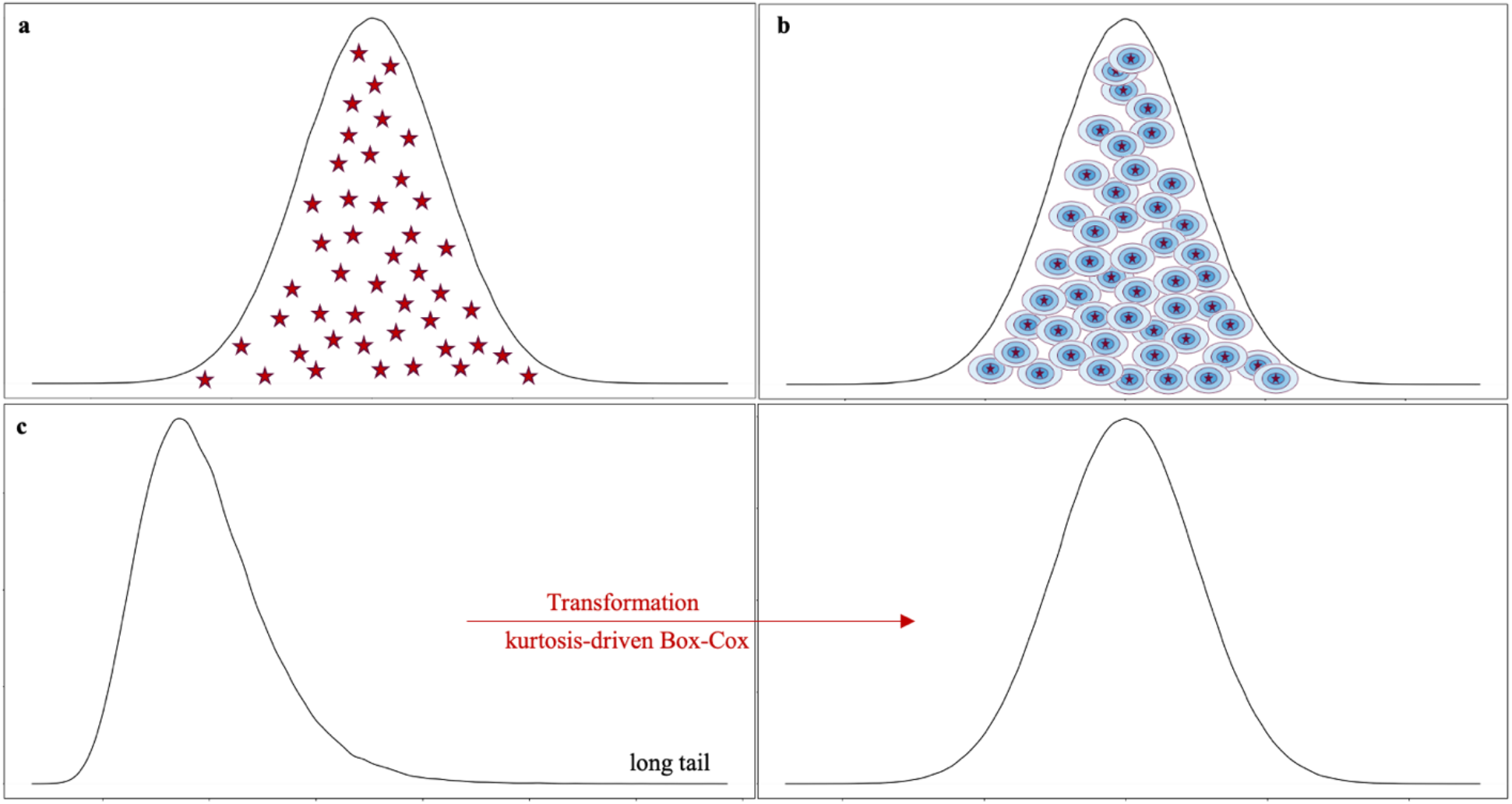
The overview of the Kernel-smoothed permutation protocol. (a) In the Naïve permutation, each permuted sample is self-representative. (b) In the Kernel-smoothed permutation, each permuted sample represents a broader region via the distribution centered by each sample. The kernel estimator learns the density as the sum of distributions placed at these samples. (c) To estimate a very small P-value in genetic association studies, the heavy-tailed distribution will reduce the effectiveness of kernel smoothing. The kurtosis-driven Box-Cox transformation relieves the problem.

### Univariate kernel-based density estimation (KDE)

Given *n* data points *X*_1_, …, *X*_*n*_ drawn from an independent and identically distributed (*iid*) sample from a population *X* with an unknown continuous probability distribution function (PDF) *f*(*x*), the kernel-based density estimator (KDE) is defined as:

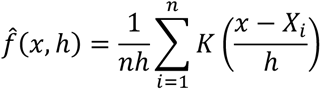

Here *K* represents the kernel function and *h* is the smoothing parameter or bandwidth. In our subsequent KDE analysis, we focused on the most used Gaussian kernel with kernel function 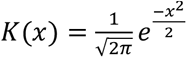.

The choice of the smoothing parameter, also known as the bandwidth *h*, is pivotal as it dictates the extent of smoothing applied. A very small value of *h* can lead to many wiggly structures on the kernel estimate, and this is a signature of under-smoothing, where the amount of smoothing is too small so some structures identified by KDE might be caused by randomness. Conversely, when *h* is too large, the computed density will be over-smoothed, where some important structures are obscured by the huge amount of smoothing, but its variance across different samples is reduced.

The performance of KDE is measured by MISE (mean integrated squared error) or AMISE (asymptotic mean integrated squared error). The core idea behind different bandwidth selection methods is to minimize the AMISE, and different methods can be viewed as different estimators to minimize the AMISE. There are several commonly used methods to select the bandwidth in KDE (Bowman, 1984; Jones et al., 1996; Scott, n.d.; Scott & Terrell, 1987; Silverman, n.d.), in our subsequent analysis, we used Silverman’s “rule of thumb” bandwidth selection method (Silverman, n.d.).

### Closed-form tests applied in our analysis

Our Kernel-Smoothed Permutation protocol is versatile and can be adapted to any distribution-free test statistics. To validate its efficacy, we applied it to three established closed-form tests: the two-sample t-test, the sequence kernel association test (SKAT) (Wu et al., 2010, 2011), and the chi-squared test.

#### Two-sample t-test

We aim to test whether there is a significant difference between two groups, A and B, within a population assuming equal variances. Under the null hypothesis, the test statistic *T* follows the studentized t-distribution and the corresponding P-value is derived as 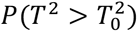.

#### Sequence kernel association test (SKAT)

SKAT is a region-based regression method designed to assess the joint effects of multiple genetic variants in a particular region on a phenotype while adjusting for covariates like principal components to account for population stratification. It employs a multiple regression model to directly regress the phenotype on variants in a region and on covariates, allowing different variants to have different directions and magnitude of effects, including no effects. This kernel association test estimates the regression coefficients of these variants using a variance-component score test within a mixed-model framework.

Supposing there are *p* variants with *n* subjects sequenced in a region. Let *y* denote the phenotype variable, *X* be the covariates matrix, and *G* stand for the genotype matrix for the *p* variants in the region. We then consider a logistic model when the phenotypes are continuous traits:

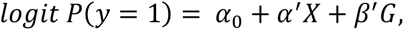

Here *α*_0_ is an intercept term, *α* is the vector of regression coefficients for the covariates, *β* is the vector of regression coefficients for the *p* observed gene variants in the region. The corresponding null hypothesis is: *H*_0_: *β* = 0. To increase the power, SKAT tests *H*_0_ by assuming each *β*_*j*_ follows an arbitrary distribution with a mean of zero and a variance of *w*_*j*_*τ*, where *τ* is a variance component and *w*_*j*_ is a pre-specified weight for variant *j*. Finally, the null hypothesis changes to *H*_0_: *τ* = 0, which can be tested using a variance-component score test: 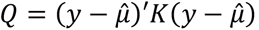,where *K* = *GWG*′, 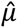 is the predicted mean of *y* under *H*_0_, *W* = *diag*(*w*_1_, …, *w*_p_) contains the weights of the *p* variants. Under the null hypothesis, *Q* follows a mixture of chi-square distributions, which can be closely approximated with the computationally efficient Davies method (Davies, 1980).

#### Chi-squared test

We aim to test for association between genotype information and binary traits (case and control) in GWAS studies using the chi-squared test. The P-value is then derived using the chi-squared distribution as 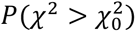.

### Data sources and quality control

For the two-sample t-test, both groups were assigned the same sample size, which was randomly selected from the set: 100, 150, 200, 250, and 500. The population mean of the first group was derived from a Uniform distribution *U*(0,1), while the population mean for the second group was fixed at 0. The population variance for the first group was chosen from the set: 1, 2, 3, 4, 5, and both groups shared this variance. Subsequently, samples for both groups were drawn from a normal distribution, using the aforementioned parameters. In R, the built-in function **t.test** facilitates the two-sample t-test and we set the significance level at 0.05.

In SKAT, the genotype information is from the representative GWAS dataset, the Wellcome Trust Case Control Consortium (WTCCC). For our analysis, we selected rheumatoid arthritis (RA) samples, comprising 4,798 individuals (1,860 cases and 2,938 controls). We excluded genetic variants with a minor allele frequency (MAF) of 1% or less, resulting in a pruned set of 392,937 variants for our subsequent analysis. The gene annotation for the human genome is from GENCODE, retaining 19,096 ENSEMBL genes for further study. The test was conducted using the **SKAT** function from the **SKAT** package in R.

For the chi-squared test, we sourced both genotype and phenotype data from the RA samples in the WTCCC, consistent with our SKAT analysis. In R, we used the built-in **chisq.test** function to execute the chi-squared test.

### Naïve permutation P-value calculation

For each round of permutation, we randomly switched labels and calculated the associated test statistic. Let *M* represent the number of permutation iterations. After *M* permutations, we obtained *X*_1_, …, *X*_*M*_ as the permuted test statistics. The P-value from the naïve permutation is given by:

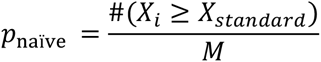

for 1 ≤ *i* ≤ *M*. Here *#*() denotes the number of times the event is satisfied. *X*_*standard*_ represents the test statistic calculated from the above closed-form tests.

In the two-sample t-test, we initially combined two groups to form a sample of *N*_*A*_ + *N*_*B*_ data points. Then we draw, without replacement, *N*_*A*_ points from the *N*_*A*_ + *N*_*B*_ points to form group A, with the remaining *N*_*B*_ points forming group B. For both the SKAT and chi-squared tests, we shuffled the “case” and “control” labels among the 4,798 RA individuals. Afterward, we computed the permuted test statistics and determined the Naïve permutation P-value for all three tests using the above equation.

### Transformations considered for permuted test statistics

Permuted test statistics derived from both the SKAT and the chi-squared test exhibit a heavy-tailed distribution, which can diminish the efficacy of KDE. To address this, we explored transforming these permuted test statistics. Our analysis considered seven distinct transformations:

1. Original transformation: retain the permuted test statistics in their original state, use *X* = (*X*_1_, …, *X*_*M*_) as permuted test statistics in kernel-smoothing.
2. Squared root transformation: use *X*^½^ as permuted test statistics in kernel-smoothing.
3. Cubic root transformation: use *X*^⅓^ as permuted test statistics in kernel-smoothing.
4. Log transformation: use ln (*X*) as permuted test statistics in kernel-smoothing.
5. Box-Cox transformation: The Box-Cox transformation is a statistical method that transforms non-normal data into an approximately normal distribution. The transformation function is given for different values of transformation parameter *λ* by the following expression.

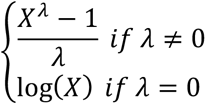 In R, we have the **boxcox** function from the **MASS** package to estimate the optimal *λ* by maximum likelihood estimation.
6. & 7. Skewness-driven and Kurtosis-driven Box-Cox transformation: While the Box-Cox transformation is designed to make data resemble a normal distribution, the results might not always exhibit the ideal skewness and kurtosis inherent to the normal distribution (a skewness of 0 and a kurtosis of 3). To address this, we introduce two additional transformations: the skewness-driven and kurtosis-driven Box-Cox transformations. The skewness-driven Box-Cox transformation optimizes the parameter λ by minimizing the absolute value of skewness in the transformed data. Conversely, the kurtosis-driven Box-Cox transformation optimizes λ to minimize the relative excess kurtosis towards a normal distribution, defined as the absolute difference between the sample kurtosis and 3. These tailored transformations allow for more nuanced adjustments, targeting specific moments of the distribution to better align with the characteristics of the normal distribution.

### Analysis procedures and settings in the Kernel-smoothed permutation protocol

Our protocol is a five-step implementation. We initiated the process with hypothesis testing for each closed-form test and obtained the corresponding P-value, which we referred to as the standard P-value. Given computational restrictions, we opted for standard P-value accuracy thresholds of 10^−7^ and 10^−8^. Next, we produced 10^8^ permuted test statistics following the procedures in *Naïve permutation P-value calculation* and calculated the Naïve permutation P-value. Third, we transformed the aforementioned 10^8^ permuted test statistics using the seven transformations previously described. We termed them as full samples. Fourth, we implemented the Kernel-smooth application (KDE) on the full samples utilizing the **kde** function from the **utilities** package in R. This function outputs a KDE object encompassing different probability functions derived from the KDE process. To compute the Kernel-smoothed permutation P-value, we employed the **pkde** function from the resulting object. To examine the effectiveness of our method under varying bandwidth lengths, we adjusted the bandwidth, calculated using Silverman’s “rule of thumb”, by multiplying it with coefficients ranging from 1 to 15 in increments of 0.5. Our objective here was to identify the optimal bandwidth coefficient and the most effective transformation, ensuring the Kernel-smoothed permutation P-value aligns closest to the standard P-value. Finally, we repeated the procedures described in the fourth step but applied them to a subset of 10^7^ transformed test statistics, which we termed 10% sub-samples.

For the two-sample t-test, our protocol was executed 500 times for two standard P-value accuracy thresholds. When performing SKAT, we repeated the above procedures 300 times for four significant genes across the two accuracy thresholds. We then derived the median from these 300 replicated P-values, designating it as the final Naïve permutation P-value and the Kernel-smoothed permutation P-value for each gene. In our exploration of the chi-squared test, each significant genetic variant across the two standard P-value thresholds was subjected to the above steps 100 times. From these iterations, we identified the median of the 100 replicated P-values, establishing it as the final Naïve permutation P-value and the final Kernel-smoothed permutation P-value for each genetic variant.

### P-value comparison between the Kernel-smoothed permutation and the Naïve permutation

To compare the effectiveness of P-values derived from the Kernel-smoothed permutation and the Naïve permutation relative to the standard P-value, we utilized the -log10(P-value) metric.

Specifically, for both Kernel-smoothed and Naïve permutation P-values, we first calculated the - log10() of the values. Subsequently, we established their deviation from the standard P-value (also expressed in -log10() terms) by taking the absolute difference. Our protocol, the Kernel-smoothed permutation, is deemed superior when its absolute -log10(P-value) difference is less than that of the Naïve permutation.

## RESULTS

### Kernel-smoothed permutation exhibits superior performance to the Naïve permutation

We first implemented our method on a two-sample t-test across two P-value accuracy thresholds based on different configurations described in **MATERIALS AND METHODS**. For a P-value accuracy threshold of 10^−8^, the Kernel-smoothed permutation outperformed Naïve permutation when an optimal bandwidth coefficient of 7 was applied, utilizing both 10^8^ original (non-transformed) permuted full samples and only 10% sub-samples (**Figure 2a**). For a P-value accuracy threshold of 10^−7^, the Kernel-smoothed permutation also outperformed Naïve permutation when an optimal bandwidth coefficient of 5 was applied, utilizing both 10^8^ original (non-transformed) permuted full samples and only 10% sub-samples (**Figure S1a**). These two optimal bandwidth coefficients are consistent with the outcomes observed in the other two closed-form tests within the same P-value accuracy threshold.

**Figure 2.**
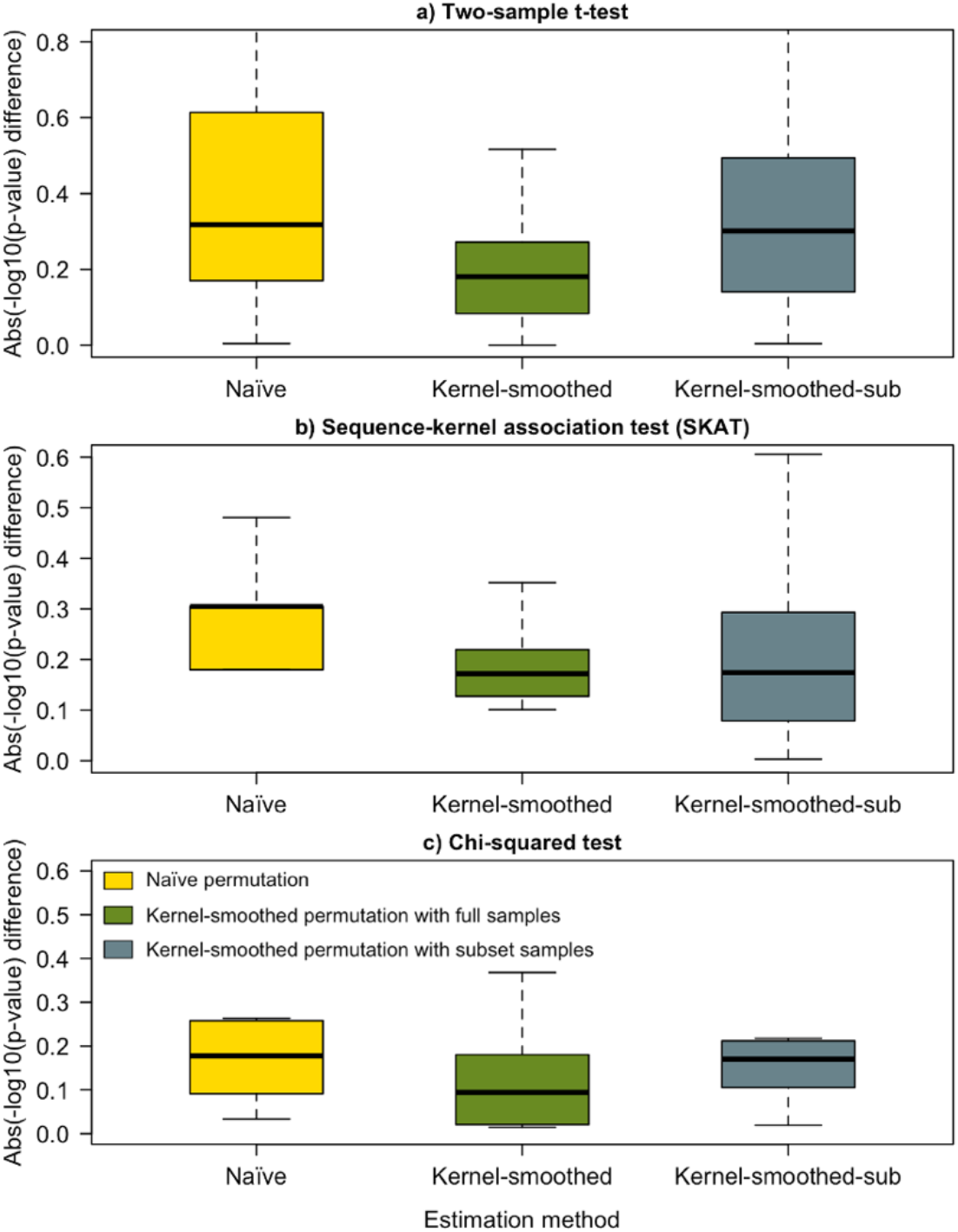
Comparison between Naïve permutation (yellow) and Kernel-smoothed permutation (olive drab for full samples and light blue for 10% sub-samples) under the optimal transformation. P-values are compared in absolute -log10(P-value) difference for a P-value accuracy threshold of 10^−8^ and an optimal bandwidth coefficient of 7. a) represents the two-sample t-test, with the optimal transformation as original (no transformation); b) represents the SKAT (gene *IP6K3*), with the optimal transformation as log transformation; and c) represents the chi-squared test, with the optimal transformation as kurtosis-driven Box-Cox transformation.

Secondly, after running SKAT on RA samples from the WTCCC dataset, we identified 4 and 15 significant genes at P-value accuracy thresholds of 10^−7^ and 10^−8^, respectively. Due to computational constraints, we restricted our application of the Kernel-smoothed permutation in SKAT to four genes across two P-value accuracy thresholds, following the configurations described in **MATERIALS AND METHODS**. Specifically, we applied Kernel-smoothed permutation to the genes *PPP1R18* and *LEMD2* for a P-value accuracy threshold of 10^−8^, and to the genes *UQCC2* and *IP6K3* for a P-value accuracy threshold of 10^−7^. In our figure illustration, we have chosen to display the results for genes *LEMD2* and *IP6K3*. However, detailed results of these four genes can be found in **Tables S1-S4**. For *IP6K3*, the Kernel-smoothed permutation outperformed Naïve permutation utilizing both 10^8^ log-transformed full samples and 10% sub-samples (**Figure 2b**). For *LEMD2*, the Kernel-smoothed permutation also outperformed the Naïve permutation utilizing both 10^8^ log-transformed full samples and 10% sub-samples but with varying results (**Figure S1b**).

Lastly, after performing the chi-squared test on RA samples from the WTCCC dataset, we identified 18 and 16 significant genetic variants at P-value accuracy thresholds of 10^−7^ and 10^−8^, respectively. We implemented our method on these significant variants across two P-value accuracy thresholds, following the configurations described in **MATERIALS AND METHODS**. For both P-value accuracy thresholds, the Kernel-smoothed permutation outperformed Naïve permutation utilizing both 10^8^ kurtosis-driven Box-Cox transformed full samples and only 10% sub-samples (**Figure 2c; Figure S1c**).

### Kurtosis-driven Box-Cox transformed samples perform the best in the Kernel-smoothed application

It is noteworthy that permuted samples derived from the two-sample t-test follow the studentized t-distribution. As the sample size increases, this t-distribution approaches the normal distribution. Given this behavior, it is justifiable to retain the permuted samples in their original state, especially since their skewness and kurtosis already closely mirror the attributes of a normal distribution.

The permuted samples generated from SKAT follow a mixture of chi-squared distribution and are heavy-tailed (**Figure 3e** and **3f; Figure S2e** and **S2f**). Employing a log transformation effectively converts these samples into a symmetric distribution characterized by optimal skewness and kurtosis aligning with a normal distribution. We also presented a P-value comparison under different transformations for both full samples and 10% sub-samples (**Figure 3a** and **3b; Figure S2a** and **S2b**). For gene *IP6K3*, log transformation performed the best among all transformations using both full samples and 10% sub-samples. For gene *LEMD2*, log transformation performed the best using full samples but performed almost the same as squared root transformation (still outperformed the others) using 10% sub-samples. When considering the full samples, we also assessed their skewness and kurtosis under different transformations (**Figure 3c** and **3d; Figure S2c** and **S2d**). The efficacy of the Kernel-smoothed permutation across various transformations seemed consistent with their respective skewness and kurtosis values. Moreover, the underlying distributions of various transformations are illustrated through their density plots, accompanied by focused visuals of the tails (**Figure 3e** and **3f; Figure S2e** and **S2f**).

**Figure 3.**
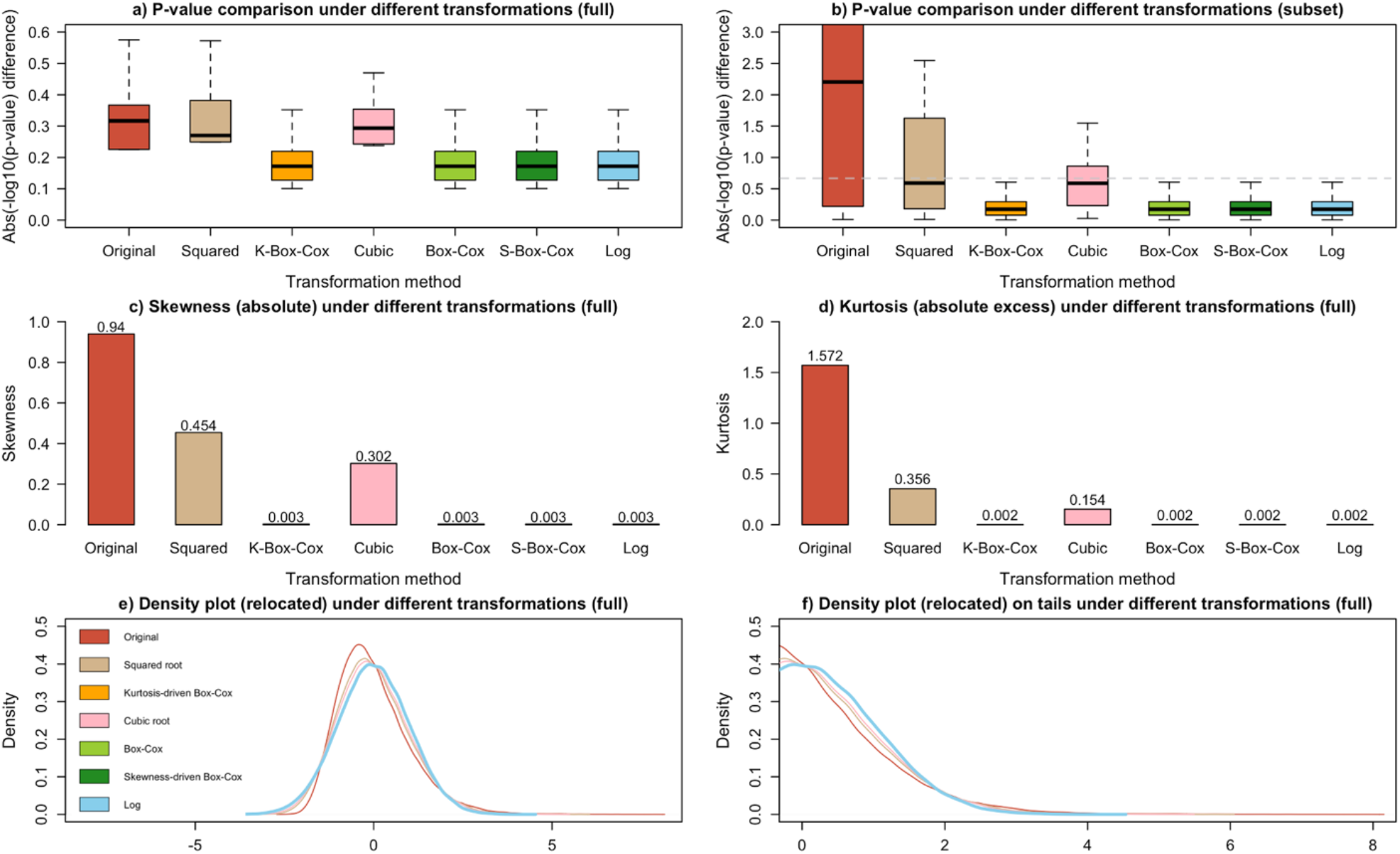
P-value comparison of Kernel-smoothed permutation under different transformations in SKAT (gene *IP6K3*). P-values are compared in absolute -log10(P-value) difference for a P-value accuracy threshold of 10^−8^ and an optimal bandwidth coefficient of 7. a) represents the P-value comparison for full samples; b) represents the P-value comparison for 10% sub-samples; c) and d) represent the skewness and kurtosis values under different transformations for full samples; and e) and f) represent the density (relocated) plots underlying distributions of various transformations along with their focused visuals of the tails for full samples.

Notably, within SKAT, the **boxcox** function estimated the optimal λ value at 0, indicating a preference for log transformation. Given that the log-transformed permuted samples already exhibit optimal skewness and kurtosis properties, there is no need to perform the three Box-Cox transformations, as they are the same in principle. Nevertheless, for the sake of visual consistency, we incorporated results under the three Box-Cox transformations in **Figure 3**, where their performance mirrored that of the log transformation.

The permuted samples generated from the chi-squared test follow a chi-squared distribution and are heavy-tailed (**Figure 4e** and **4f; Figure S3e** and **S3f**). Employing a Kurtosis-driven Box-Cox transformation effectively converts these samples into a symmetric distribution characterized by optimal kurtosis aligning with a normal distribution. We also presented a P-value comparison under different transformations for both full samples and 10% sub-samples (**Figure 4a** and **4b; Figure S3a** and **S3b**). For a P-value accuracy threshold of 10^−8^, Kurtosis-driven Box-Cox transformation performed the best among all transformations using full samples but with varying results compared to the squared root transformation but performed the best using 10% sub-samples. For a P-value accuracy threshold of 10^−7^, Kurtosis-driven Box-Cox transformation did not outperform the squared root transformation but still outperformed the others using full and 10% sub-samples. Remarkably, during the chi-squared test, the **boxcox** function determined that the optimal λ value for the Kurtosis-driven Box-Cox transformation was approximately 0.45. This value is quite close to the transformation parameter of 0.5 typically employed in the square root transformation. Consequently, this similarity in transformation parameters partly accounts for the occasional lack of significant improvement observed with the Kurtosis-driven Box-Cox transformation when compared to the square root transformation. When considering the full samples, we also assessed their skewness and kurtosis under different transformations (**Figure 4c** and **4d; Figure S3c** and **S3d**). The efficacy of the Kernel-smoothed permutation across various transformations seemed almost consistent with their respective skewness and kurtosis values. Moreover, the underlying distributions of various transformations are illustrated through their density plots, accompanied by focused visuals of the tails (**Figure 4e** and **4f; Figure S3e** and **S3f**). Detailed information on the above results can be found in **Tables S5-S12**.

**Figure 4.**
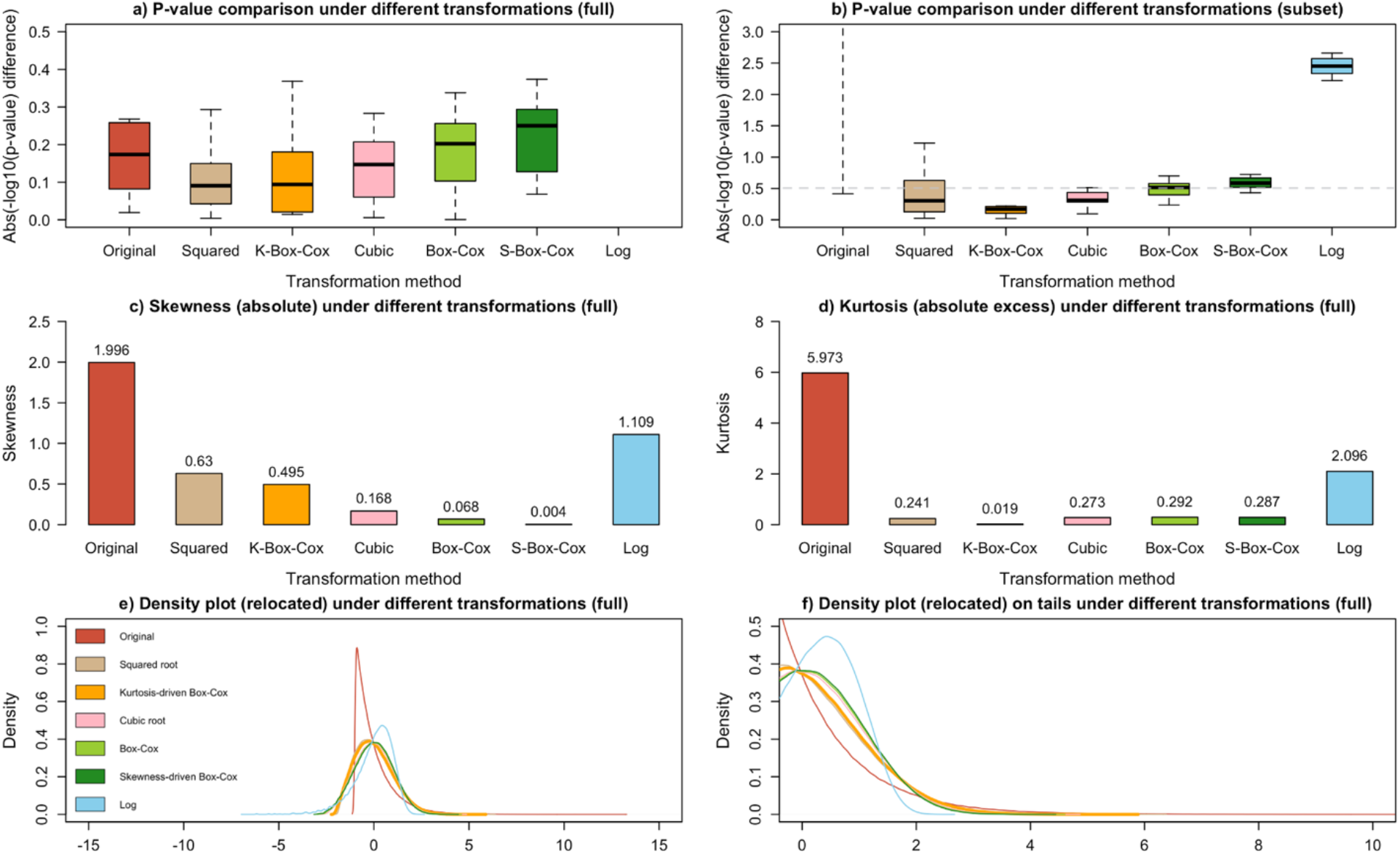
P-value comparison of Kernel-smoothed permutation under different transformations in the chi-squared test. P-values are compared in absolute -log10(P-value) difference for a P-value accuracy threshold of 10^−8^ and an optimal bandwidth coefficient of 7. a) represents the P-value comparison for full samples; b) represents the P-value comparison for 10% sub-samples; c) and d) represent the skewness and kurtosis values under different transformations for full samples; and e) and f) represent the density (relocated) plots underlying distributions of various transformations along with their focused visuals of the tails for full samples.

## DISCUSSION

In conclusion, we introduced the Kernel-smoothed permutation, a technique that utilizes kurtosis-driven Box-Cox transformation-guided KDE to model the null distribution of permuted test statistics, enhancing the precision of small P-value calculations. Our approach was tested across three closed-form tests, and consistently demonstrated superiority over the Naïve permutation in terms of P-value accuracy and efficiency in both full and 10% sub-samples, reducing the number of permutations needed. As a non-parametric strategy, the Kernel-smoothed permutation does not hinge on any parametric prerequisites and can be seamlessly incorporated with any distribution-free test statistics. We anticipate that our methodology will be similarly effective for a broader spectrum of test statistics.

Our study presents certain limitations that merit consideration. Firstly, our validation of the Kernel-smoothed permutation protocol focuses on its application to three predefined closed-form tests. Nonetheless, our methodology has the potential to be extended to test statistics with indeterminate distributions, such as those utilized in the interaction-integrated linear mixed model (ILMM).(Li et al., 2023). Secondly, our current transformation approaches primarily target the 3rd (skewness) and 4th (kurtosis) moments. Future endeavors could involve transformations by higher-order moments and explore their relative efficacy in estimating small P-values. Furthermore, we have yet to provide closed-form mathematical derivations that theoretically quantify the superiority of the Kernel-Smoothed Permutation. Lastly, the potential of the Kernel-smoothed permutation framework extends beyond our current scope, encompassing a broader range of omics data and testing techniques, making it a versatile tool in genetic studies for approximating small P-values. Exploration of these expanded applications remains a subject for forthcoming research.

## AUTHOR CONTRIBUTIONS

Convinced the study: J.B., J.W., Q.L. Implemented the tool: J.B., J.W. Analyzed data: J.B., C.L., J.W., Q.L. Supervised the study: J.W., Q.L. Wrote the manuscript: A.B., Q.L. with contributions from all authors.

## ACKNOWLEDGMENTS

This work is supported by the New Frontiers in Research Fund [Exploration NFRFE-2018-00748 to Quan Long], the Alberta Innovates LevMax-Health Program Bridge Funds [222300769 to Quan Long], the Canada Foundation for Innovation [36605 to Quan Long], NSERC Discovery Grant [RGPIN-2018-04328 to Jingjing Wu and RGPIN-2017-04860 to Quan Long], and the Alberta Innovates Graduate Student Scholarships to Jiayi Bian.

## CONFLICT OF INTEREST STATEMENT

The authors declare no conflict of interest.

## DATA AVAILABILITY STATEMENT

The gene model file used in SKAT is openly available at https://www.gtexportal.org/home/datasets/gencode.v26.GRCh38.genes.gtf. The WTCCC dataset is available at https://www.wtccc.org.uk/. The SKAT package is available at https://cran.r-project.org/web/packages/SKAT/SKAT.pdf; the Utilities package is available at https://cran.r-project.org/web/packages/utilities/index.html; and the MASS package is available at https://cran.r-project.org/web/packages/MASS/index.html. Bandwidth calculation methods in KDE is available at https://www.rdocumentation.org/packages/stats/versions/3.6.2/topics/bandwidth. Kernel-smoothed permutation is freely available on GitHub under the MIT license and can be found at https://github.com/theLongLab/Kernel_smoothed_permutation.

## Notes

### Competing Interest Statement

The authors have declared no competing interest.

