## Supplementary Figures for "Kernel-smoothed permutation for extreme P-value estimation in genetic association studies"

**Figure S1** Comparison between Naïve permutation (yellow) and Kernel-smoothed permutation (olive drab for full samples and light blue for 10% sub-samples) under the optimal transformation. P-values are compared in absolute -log10(p-value) difference for a p-value accuracy threshold of ${10}^{-7}$ and an optimal bandwidth coefficient of 5. a) represents the two-sample t-test, with the optimal transformation as original (no transformation); b) represents the SKAT (gene *LEMD2*), with the optimal transformation as log transformation; and c) represents the chi-squared test, with the optimal transformation as kurtosis-driven Box-Cox transformation.

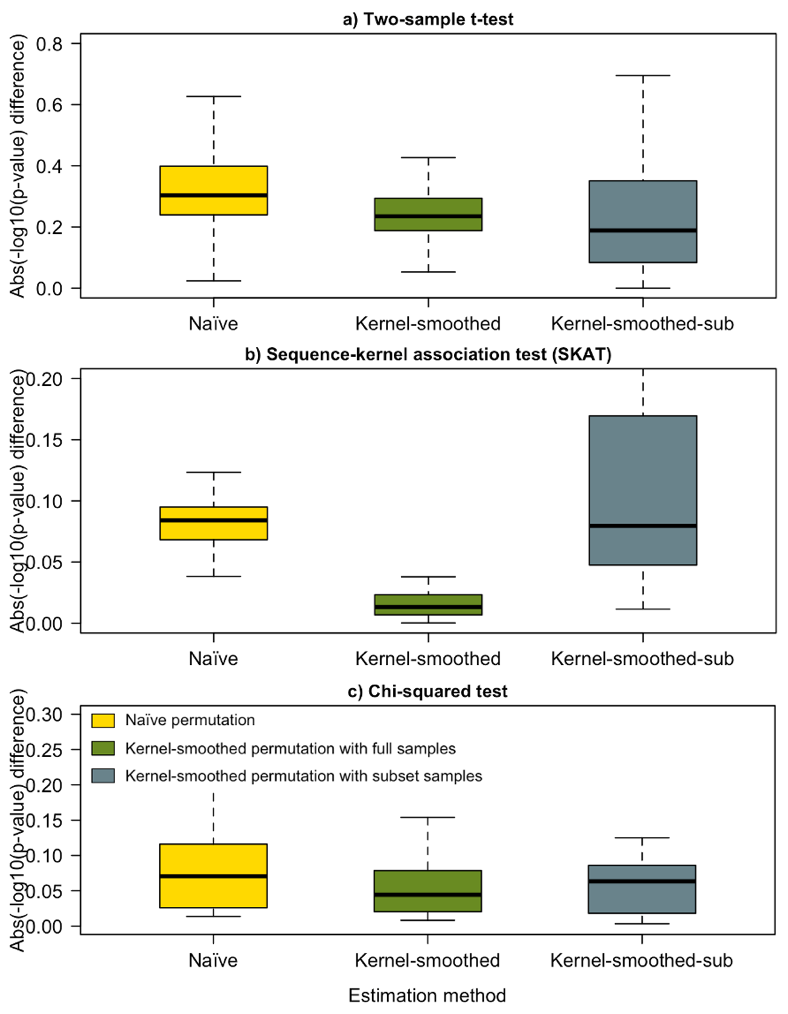

**Figure S2** P-value comparison of Kernel-smoothed permutation under different transformations in SKAT (gene *LEMD2*). P-values are compared in absolute -log10(p-value) difference for a p-value accuracy threshold of ${10}^{-7}$ and an optimal bandwidth coefficient of 5. a) represents the p-value comparison for full samples; b) represents the p-value comparison for 10% sub-samples; c) and d) represent the skewness and kurtosis values under different transformations for full samples; and e) and f) represent the density (relocated) plots underlying distributions of various transformations along with their focused visuals of the tails for full samples.

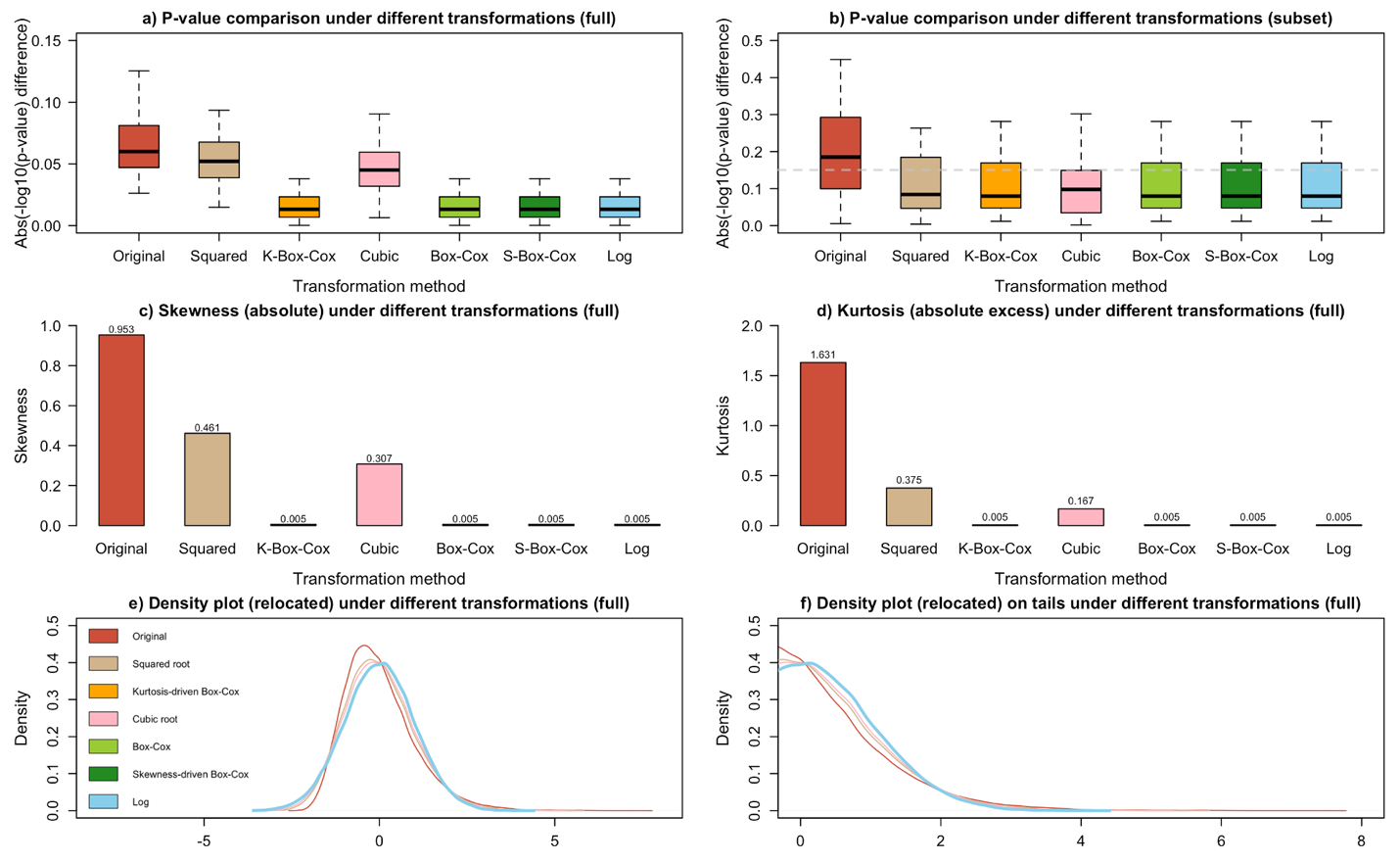

**Figure S3** P-value comparison of Kernel-smoothed permutation under different transformations in the chi-squared test. P-values are compared in absolute -log10(p-value) difference for a p-value accuracy threshold of ${10}^{-7}$ and an optimal bandwidth coefficient of 5. a) represents the p-value comparison for full samples; b) represents the p-value comparison for 10% sub-samples; c) and d) represent the skewness and kurtosis values under different transformations for full samples; and e) and f) represent the density (relocated) plots underlying distributions of various transformations along with their focused visuals of the tails for full samples.

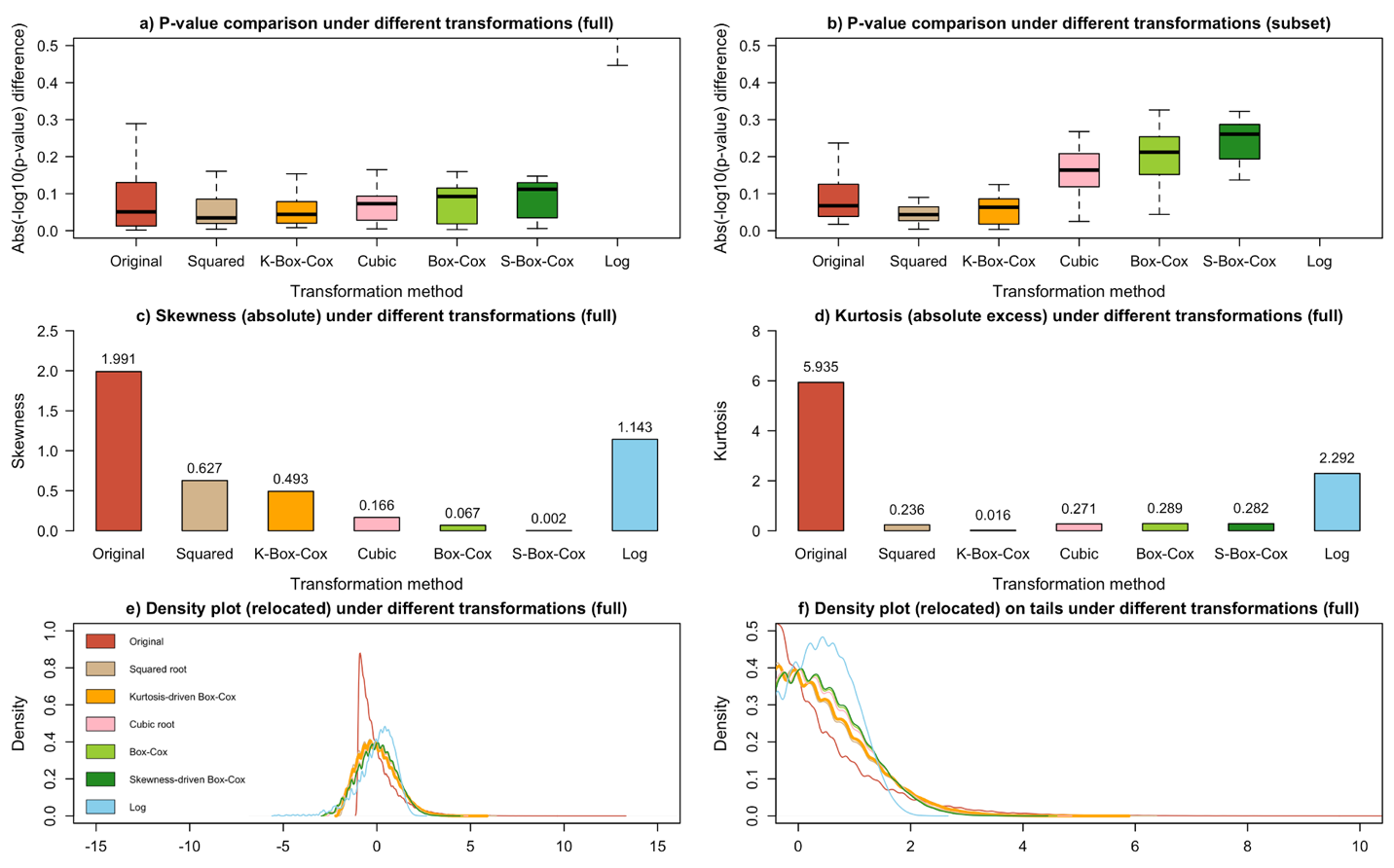
